# LINC02428, a liver-specific long noncoding RNA suppresses hepatocellular carcinoma progression

**DOI:** 10.1101/2022.09.21.508837

**Authors:** Qiangnu Zhang, Lesen Yan, Jiaojuan Chen, Liping Liu

**Author notes:** Correspondence, Liping Liu, Division of Hepatobiliary and Pancreas Surgery, Department of General Surgery, Shenzhen People’s Hospital (The Second Clinical Medical College, Jinan University; The First Affiliated Hospital, Southern University of Science and Technology), 1017 Dongmen Bei Lu, 518020 Shenzhen, China,.

## Abstract

There is growing evidence that lncRNAs play an important role in the progression of HCC, especially liver-specific expression of lncRNAs. LINC02428 is a liver-specific labeled lncRNA, but its role in HCC is unclear. We investigated the expression characteristics of LINC02428 in hepatocellular carcinoma tissues and analyzed its function in HCC. Using TCGA public data we found that the expression level of LINC02428 was significantly reduced in hepatocellular carcinoma tissues, compared to normal tissues. LINC02428 was also significantly reduced in hepatocellular carcinoma tissues in our cDNA microarray data from HCC patients. Survival analysis suggests that HCC patients with lower LINC02428 have a lower overall survival rate. In an in vitro analysis, overexpression of LINC02428 inhibited HCC cell proliferation, migration, and induced apoptosis. In conclusion, we report for the first time the expression profile of a liver-specific lncRNA, LINC02428, in HCC and confirmed its ability to inhibit the progression of HCC. LINC02428 may become a new target for the diagnosis and treatment of HCC.

## Introduction

In 2020, there were approximately 0.9 million new cases of liver cancer and 0.8 million deaths due to liver cancer worldwide, which were the sixth highest number of cases of cancer and the third highest number of deaths from cancer worldwide, respectively^[1]^. Hepatocellular carcinoma (HCC) accounts for 85–95% of primary liver cancer. Approximately 80% of HCC patients are in the advanced stages when diagnosed, thus losing the opportunity for surgery. The overall 5-year survival rate is less than 30%, with an 80% recurrence rate in advanced HCC patients^[2, 3]^. Therefore, explaining the mechanism of HCC and developing effective treatment measures are urgent needs.

Long noncoding RNAs (lncRNAs), with a length of >200 nucleotides, play their roles by functioning as signals, miRNA sponges, decoys, guides, or scaffolds for other regulatory proteins^[4, 5]^. Accumulating evidence has suggested that lncRNAs are involved in HCC and regulate HCC proliferation, migration, invasion, and therapy resistance^[6–8]^. Some lncRNAs are tissue-specific, and lncRNAs expressed specifically in the liver may be more important for the development of hepatocellular carcinoma in the liver than non-tissue-specific lncRNAs, Hence the liver-specific expression of lncRNAs needs to be emphasized. For instance, He’s team identified a novel, liver-specific long noncoding RNA LINC01093 that suppresses HCC progression^[9]^. Mo reported that FAM99B is a liver-specific lncRNA and suppresses HCC progression^[10]^. We identified a liver-specific expressed lncRNA, LINC02428, whose expression characteristics and function in hepatocellular carcinoma have not been reported. In this study, we compared the difference in expression of LINC02428 in HCC and peri-cancerous normal tissues and analyzed its effect on the proliferation and migration of HCC.

## Materials and Methods

### Public data collection

The RNA sequencing (RNA-seq) data and clinical data of The Cancer Genome Atlas project (TCGA-LIHC) were collected from UCSC Xena public datahub (http://xena.ucsc.edu/public/). The gene tissue specificity index of lncRNAs were obtained from Human Ageing Genomic Resources (https://genomics.senescence.info/gene_expression/tau.html). The Pan-cancer data of LINC02428 were obtained from GEPIA database (http://gepia.cancer-pku.cn/).

### HCC specimens and cell lines

cDNA microarray data of 90 HCC patients (HLivH090Su01) were obtained from the Shanghai Engineering Center for Molecular Medicine. HUH7, PLC/PRF/5 were obtained from the Cell Bank of the Chinese Academy of Sciences. All cell lines were cultured in Dulbecco’s modified Eagle’s medium (DMEM, Gibco, CA, USA) with 10% fetal bovine and 1% penicillin/streptomycin in 5% CO2 at 37°C.

### Cell viability assay

Transfected or treated cells were planted into a 96-wells plate at a density of 3000 cells/100 μL. CellTiter-Lumi™ Steady Plus Luminescent Cell Viability Assay Kit was used to detect the cell s viability changes at multiple time points. 100 μL detection reagent was added for each well. The luminescence intensity was detected by a luminometer (SPARK 10M) after 10 min incubation.

### Colony formation

The indicated number of cells were planted in a 6-well plate and cultured in an incubator for 3-10 days. Colonies c were stained with 0.1% crystal violet dye and subsequently counted.

### Healing assay assay

For wound healing assay, 2-Well ibidi Culture-Inserts were used and 70 μL cells (30 0000cells/mL) were planted for each well. The gap was made after 24 h and images were obtained at indicated time point.

### Analysis of apoptosis

The apoptosis of treated cells was continuously monitored by the RealTime Glo Annexin V Apoptosis Assay kit. In brief, 5000 cells were planted into a 96-well plate in 100 μL DMEM. 0.1 μL Annexin V NanoBiT™ Substrate, 0.1 μL Annexin V-SmBiT, and 0.1 μL Annexin V-LgBiT were added to each well. Luminescence was read by a luminometer (SPARK 10M).

### Statistical analysis

R 4.1.0 software (https://www.R-project.org/) and GraphPad Prism 7.0 (GraphPad Software, CA, USA) was used for statistical analysis. Data are presented as mean ± standard deviation. Differences between two different groups were evaluated using a t-test. Differences between two different groups in the RNA-seq data or microarray data were evaluated using the Willcox test. Pearson’s correlation or Spearmans correlation analyses were used to analyzing correlations. Survival data were analyzed using the Kaplan–Meier method or Cox proportional hazards models. Classified variables were analyzed using the Chi-squared test. *P* < 0.05 was defined as statistical significance.

## Results

### LINC02428 is a liver-specific long noncoding RNA that reduced in HCC

We used Palmers algorithm^[11]^, Tau Index of Gene Tissue Specificity, to assess the specificity of LINC02428 expression among different tissues. The results revealed that LINC02428 was specifically expressed in the liver (Figure 1A). In the TCGA public dataset, LINC02428 was only expressed in hepatocellular carcinoma and cholangiocarcinoma, and endogenous expression of LINC02428 was barely detectable in other cancer types. In the TCGA public dataset, LINC02428 was only expressed in hepatocellular carcinoma and cholangiocarcinoma, and endogenous expression of LINC02428 was barely detectable in other cancer types (Figure 1B). In the TCGA-LIHC data and the GTEx data, LINC02428 was significantly low expressed in liver cancer tissues (P<0.05), compared with normal tissues (Figure 1C). The TCGA-LIHC data also showed that LINC02428 levels further decreased with the progression of TNM staging (Figure 1D). In our collection of cDNA microarray data from liver cancer patients, we confirmed the low expression of LINC02428 in liver cancer tissues using qRT-PCR (Figure 1E). Also in fresh tissues from liver cancer patients, we verified the low expression of LINC02428 using FISH probes (Figure 1F).

**Figure 1.**
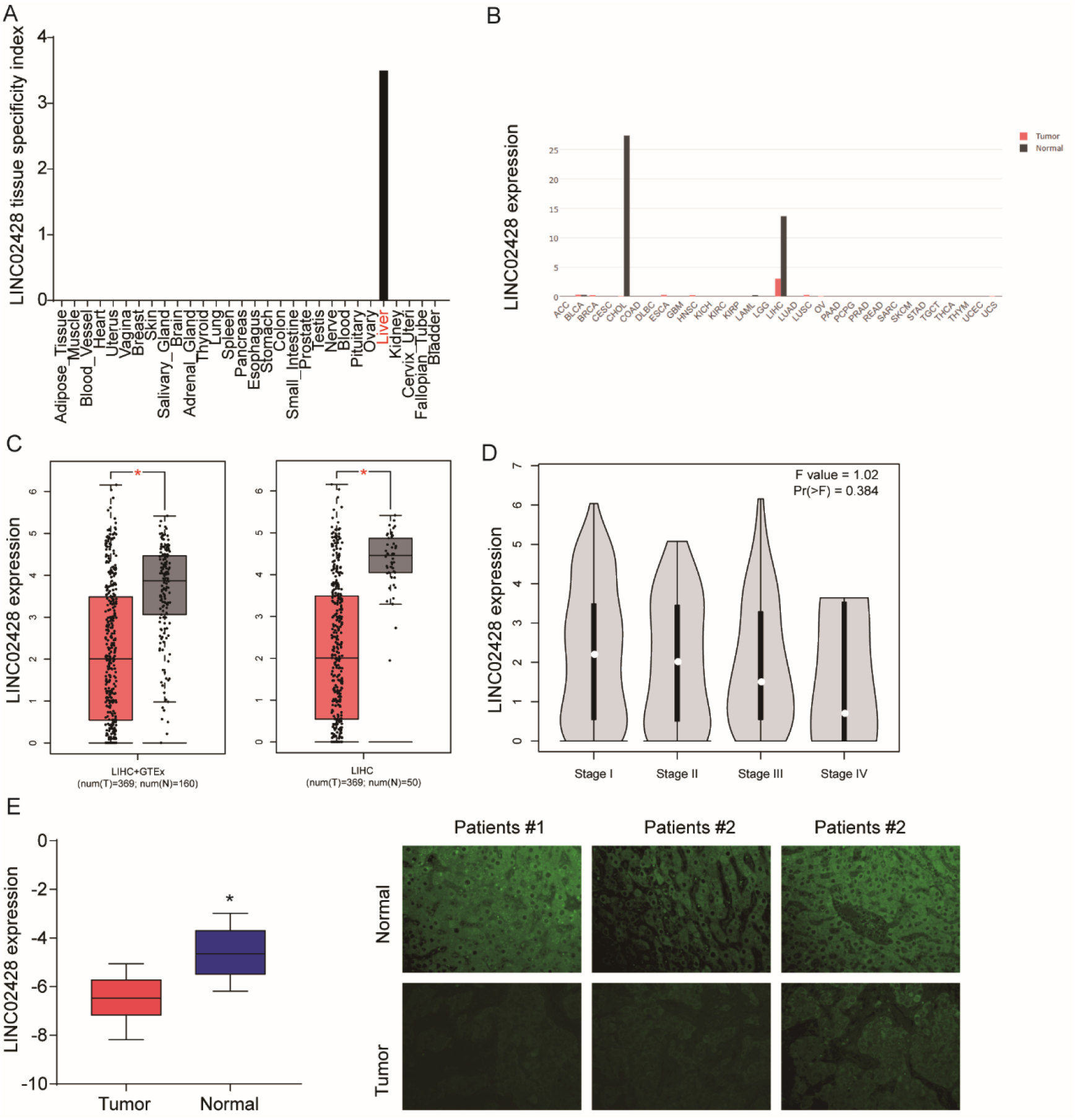
LINC02428 is a liver-specific long noncoding RNA that is reduced in HCC. (A) Tau Index of Gene Tissue Specificity for LINC02428 among different tissues; (B) LINC02428 level in different cancer types. Data were obtained from TCGA; (C) LINC02428 level in HCC and normal tissues;(D) qRT-PCR detected the LINC02428 level changes in HCC and normal tissues. cDNA collected from 64 HCC patients; (E) FISH probe detected LINC02428 levels in fresh HCC tissues.

### Low expression of LINC02428 is associated with poor prognosis in patients with HCC

We divided patients from TCGA-LIHC into a high LINC02428 group and a low LINC02428 group using the median value of LINC02428 as the cut-off value. k-M survival analysis found that patients in the low LINC02428 group had poorer overall survival (Figure 2A), but LINC02428 levels were not associated with disease-free survival (Figure 2B). We also verified the relationship between LINC02428 and overall survival using cDNA microarray data from HCC patients(Figure 2C). These data suggest that low LINC02428 is associated with poor patient prognosis

**Figure 2.**
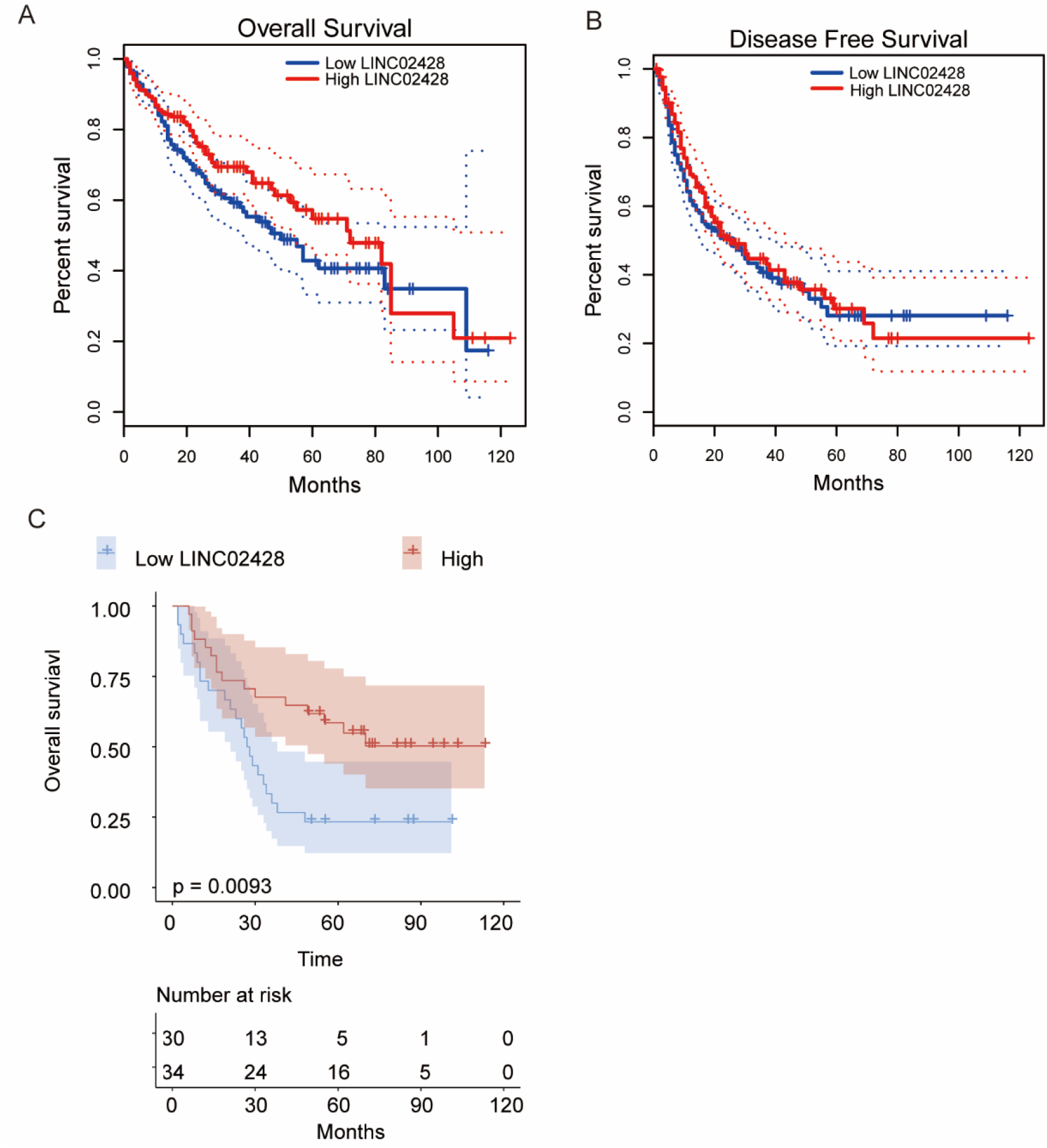
Low expression of LINC02428 associated with poor prognosis of patients with HCC.(A-B) Patients in TCGA-LIHC were divided into a high LINC02428 group and a low LINC02428 group using the median value of LINC02428 as the cut-off value. k-M survival analyses were performed to estimate the association of the LINC02428 level with overall survival and disease-free survival;(C) The association of the LINC02428 level with overall survival in 64 HCC patients.

### LINC02428 suppresses proliferation and induced apoptosis of HCC cells

Overexpression of LINC02428 significantly reduced the cellular activity of HUH7 and PLC/PRF/5 cells (Figure 3A). Clonogenic assay data suggested that overexpression of LINC02428 could inhibit the proliferation of HCC cells (Figure 3B). Real-time regulatory death analysis revealed that overexpression of LINC02428 could induce regulatory death in HCC cells (Figure 3C)

**Figure 3.**
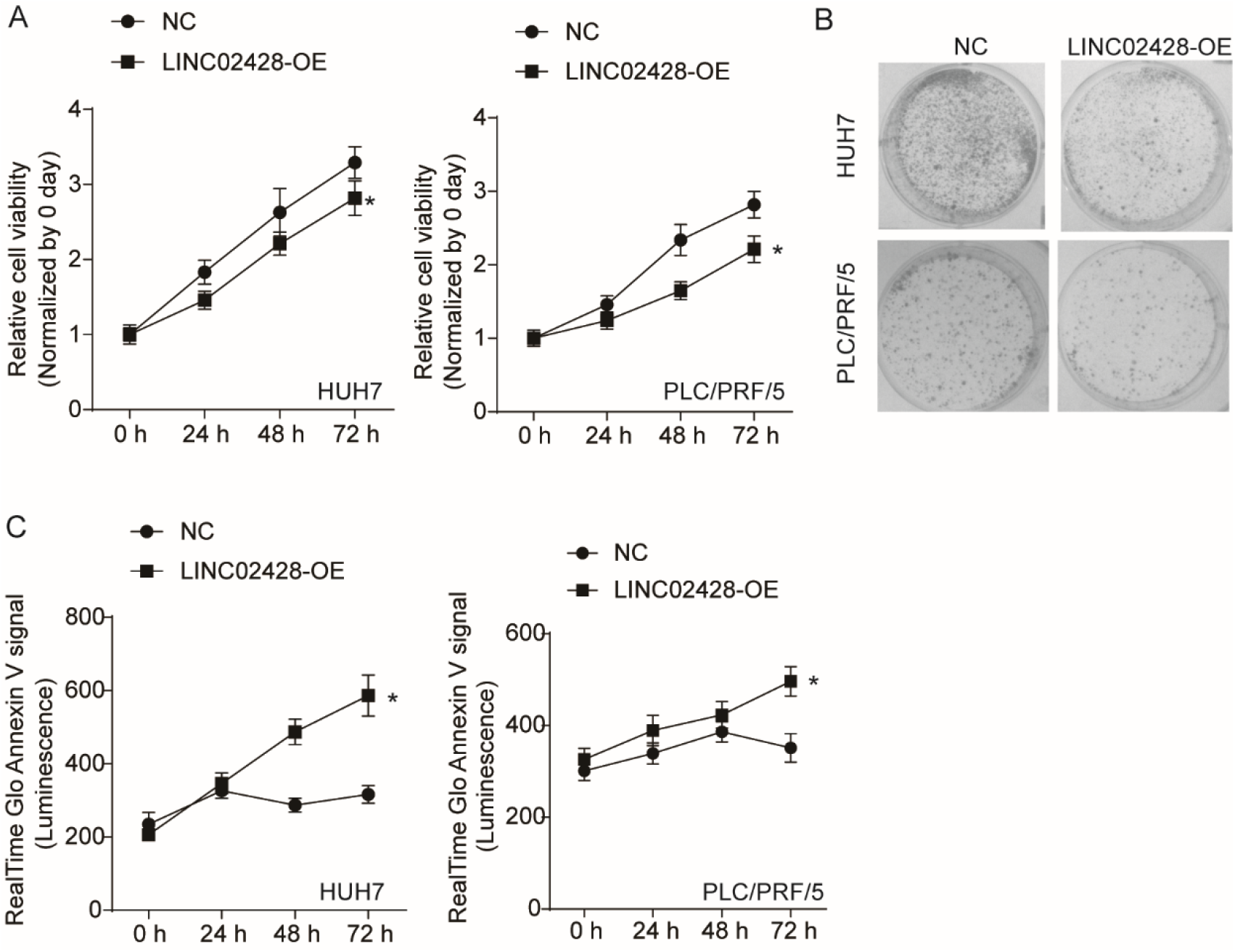
LINC02428 suppresses proliferation and induced apoptosis of HCC cells. (A)CCK-8 assay was used to estimate the cell viability changes after LINC02428 overexpression; (B) Clone formation assay for HUIH7 and PLC/PRF/5 cells with LINC02428 overexpression; (C)Real-time apoptosis assay for HUIH7 and PLC/PRF/5 cells with LINC02428 overexpression; *P<0.05.

### LINC02428 inhibited the migration of HCC cells

Transwell analysis suggested that overexpression of LINC02428 could inhibit the migration of HCC cells (Figure 4A). Similar results were obtained by scratch assay (Figure 4B).

**Figure 4.**
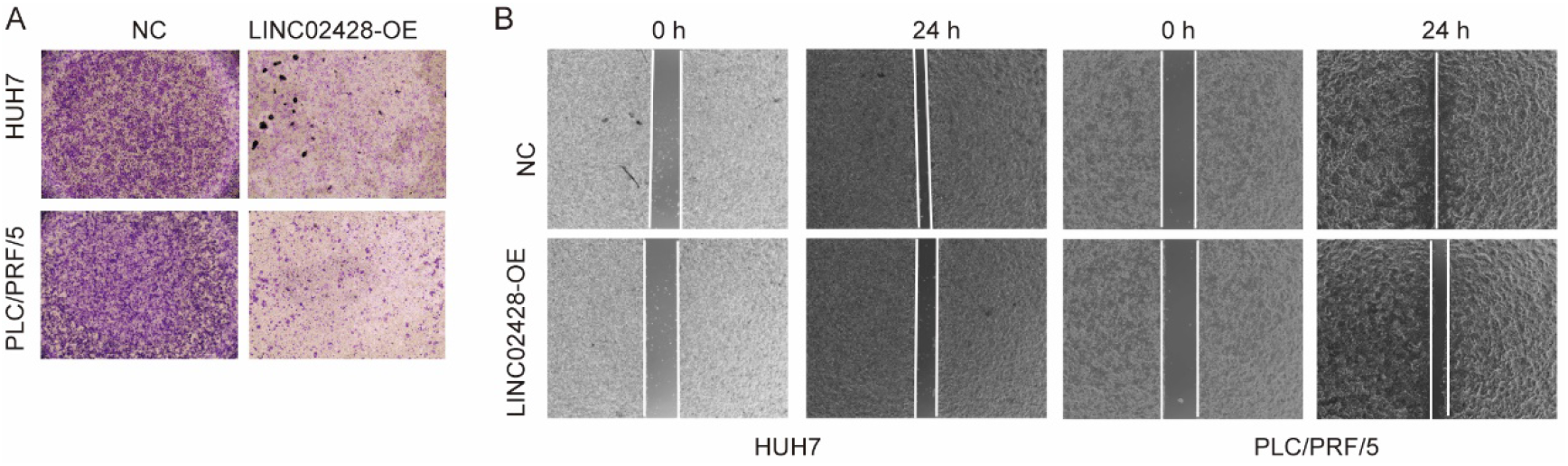
LINC02428 inhibited migration of HCC cells. (A)Transwell assay for HUIH7 and PLC/PRF/5 cells with LINC02428 overexpression;(B)Wound healing assay HUIH7 and PLC/PRF/5 cells with LINC02428 overexpression.

## Discussion

LncRNAs are involved in various pathophysiological processes including cell growth, apoptosis, and metastasis in HCC^[12]^. In the present study, we identified LINC02428 as a liver-specific expressed lncRNA, a knowledge that was reported for the first time. Using public data from TCGA we found that LINC02428 was abundantly expressed endogenously only in HCC and cholangiocarcinoma, while it was hardly detected in other cancers. Further expression analysis we found that LINC02428 was present in HCC with low expression, compared to normal tissues of cancer. This expression profile was validated in external cDNA microarray data. Survival analysis suggests that HCC patients with lower LINC02428 levels have a poorer prognosis. In vitro cellular assays also confirmed that overexpression of LINC02428 inhibited the progression of HCC. These data suggest a role for LINC02428 as a tumor suppressive gene in HCC.

Liver-specific lncRNAs are more closely related to the development of HCC and are more suitable as biomarkers and therapeutic targets for liver cancer. Several liver-specific lncRNAs that play important functions in HCC have now been reported. Zheng et al. reported that LINC01554 induced glucose metabolism reprogramming by reduced PKM2 expression in HCC. These signaling changes suppressed the tumorigenicity of HCC^[13]^. Ding et al. also demonstrated that a low level of LINC01554 inhibited HCC^[14]^. LINC02499 is another reported liver-specific long non-coding RNA. It could be a potential novel diagnostic and prognostic biomarker for HCC^[15]^. Zhang et al. found that LINC00261 suppresses cell proliferation, invasion, and Notch signaling pathways in HCC^[16]^. And EZH2 induces epigenetic regulation that may reduce LINC00261 in HCC^[17]^. Another liver-specific LINC01146 was found could inhibit the malignant phenotype of HCC cells both in vitro and in vivo, hence regarded as a promising prognostic marker^[18]^. Li reported LINC02362 attenuates HCC progression through sponging the miR-516b-5p^[19]^. These reported liver-specific lncRNAs as well as our reported LINC02428 have greater potential for clinical application. Follow-up studies should focus on how these lncRNAs can be used as diagnostic markers or therapeutic targets.

lncRNAs perform biological functions in four main modes, including signal, decoy, guide, and scaffold^[20, 21]^. Recent studies have shown that although lncRNAs cannot encode proteins, some lncRNAs have the ability to encode peptides^[21, 22]^. These lncRNAs also encode peptides with important roles. The mechanism of LINC02428 has not been investigated in this study yet, but we found that there is a small ORF region with encoding potential in LINC02428. We will focus on this key point in future studies.

In conclusion, we report for the first time the expression characteristics and functions of liver-specific lncRNA, LINC02428, in HCC. LINC02428 may become a new target for liver cancer therapy.

